# Frequent *PIK3CA* mutations in eutopic endometrium of patients with ovarian clear cell carcinoma

**DOI:** 10.1101/2021.02.25.432943

**Authors:** Kosuke Murakami, Akiko Kanto, Kazuko Sakai, Chiho Miyagawa, Hisamitsu Takaya, Hidekatsu Nakai, Yasushi Kotani, Kazuto Nishio, Noriomi Matsumura

## Abstract

Recent studies have reported cancer-associated mutations in normal endometrium. Mutations in eutopic endometrium may lead to endometriosis and endometriosis-associated ovarian cancer. We investigated *PIK3CA* mutations (*PIK3CA*m) for three hotspots (E542K, E545K, H1047R) in eutopic endometrium in patients with ovarian cancer and endometriosis from formalin-fixed paraffin-embedded specimens by laser-capture microdissection and droplet digital PCR. The presence of *PIK3CA*m in eutopic endometrial glands with mutant allele frequency ≥15% were as follows: ovarian clear cell carcinoma (OCCC) with *PIK3CA*m in tumors, 20/300 hotspots in 11/14 cases; OCCC without *PIK3CA*m, 42/78 hotspots in 11/12 cases; high-grade serous ovarian carcinoma, 8/45 hotspots in 3/5 cases; and endometriotic cysts, 5/63 hotspots in 5/6 cases. These rates were more frequent than in non-cancer non-endometriosis controls (7/309 hotspots in 5/17 cases). In OCCC without *PIK3CA*m, 7/12 (58%) cases showed multiple hotspot mutations in the same eutopic endometrial glands. In 3/54 (5.6%) cases, *PIK3CA*m was found in eutopic endometrial stroma. Multi-sampling of the OCCC tumors with *PIK3CA*m showed intratumor heterogeneity in three of eight cases. In two cases, *PIK3CA*m was detected in the stromal component of the tumor. Homogenous *PIK3CA*m in the epithelial component of the tumor matched the mutation in eutopic endometrial glands in only one case. Eutopic endometrial glands in ovarian cancer and endometriosis show high frequency of *PIK3CA*m that is not consistent with tumors, and multiple hotspot mutations are often found in the same glands. Our results suggest that most *PIK3CA*m in eutopic endometrial glands are passenger rather than driver mutations.

## Introduction

Ovarian cancer is a gynecologic malignancy with one of the worst prognoses. Ovarian clear cell carcinoma (OCCC) occurs frequently in Japan and accounts for approximately one quarter of all ovarian cancer cases [1]. OCCC is an endometriosis-associated ovarian cancer (EAOC) [2] and was previously thought to arise from endometriotic cysts (ECs) [3]. However, later studies have not confirmed this possibility, and the origin of OCCC remains unclear.

Recent advances in sequencing technology have revealed the presence of numerous cancer-associated mutations in the eutopic endometrial epithelium in healthy patients [4,5]. Cancer-associated mutations, including mutations in *PIK3CA*, are frequently found in deep infiltrating endometriosis (DIE) [6]. Because the same mutations in eutopic endometrium are also found in ECs, endometriosis is thought to be caused by the reflux of eutopic endometrium with gene mutations [7]. Notably, a case of clonal lineage from eutopic endometrium to endometriosis and OCCC was reported [8]. Therefore, eutopic endometrium has recently attracted a great deal of interest as a potential origin of EAOC [9]. A previous study showed that endometrial glands are composed of monoclonal cell populations [10]. Therefore, to clarify whether OCCC arises from eutopic endometrial cells, it is necessary to perform mutation analysis for individual endometrial glands and compare the results with mutations in ovarian tumors.

Droplet digital PCR (ddPCR) is a technology that amplifies even a very small amount of DNA to enable mutation analysis [11]. In a recent study in endosalpingiosis, the authors collected formalin-fixed paraffin-embedded (FFPE) samples from glands by laser capture microdissection (LCM), and ddPCR was used to analyze hotspot mutations in *BRAF* and *KRAS* [12]. We hypothesized that this technique would allow for mutation analysis of individual endometrial glands from FFPE samples of the uterus.

Specific mutations in *PIK3CA* have been shown to activate the PI3K/Akt/mTOR pathway and are deeply involved in human carcinogenesis [13]. *PIK3CA* is one of the most frequently mutated genes along with *ARID1A* in OCCC [14], and co-mutation of *Pik3ca* and *Arid1a* in mouse ovaries causes cancer similar to human OCCC [15]. In human cancers, including OCCC, three hotspot mutations in *PIK3CA* have been identified: E542K, E545K, and H1047R [13,16,17].

In this study, we examined *PIK3CA* hotspot mutations in eutopic endometrium in ovarian cancer, especially OCCC, and endometriosis cases by ddPCR and compared them with mutations in tumors. This study is the first report to compare mutations between tumors and normal eutopic endometrium in multiple ovarian cancer cases.

## Materials and Methods

### Sample collection

Patients who underwent surgery at the Department of Obstetrics and Gynecology, Kindai University Hospital between January 1999 and December 2019 were included in this study. For OCCC cases, FFPE samples of tumors of the ovary, eutopic endometrium, and co-existing endometriosis were obtained. We also obtained FFPE samples of tumors of the ovary and eutopic endometrium for high-grade serous ovarian carcinoma (HGSOC) and EC cases. As controls, FFPE samples of eutopic endometrium were obtained from patients whose uterus had been removed due to benign disease or cervical intraepithelial neoplasia without endometriosis or adenomyosis. Pathological diagnosis of each sample was made by two pathologists. This study was conducted with the approval of the Institutional Review Board of Kindai University Faculty of Medicine (27-182). Patients in this study were given an appropriate opportunity to refuse to participate in the study by opt out on the website of Kindai University Faculty of Medicine (https://www.kindai.ac.jp/medicine/).

### DNA extraction

For OCCC and HGSOC tumors, after confirming the tumor area by hematoxylin and eosin staining, FFPE samples were sliced at a thickness of 3 μm and macrodissection was performed using a razor to obtain only the tumor area. The collected samples were deparaffinized, and DNA was extracted using the QIAamp DNA Micro Kit (QIAGEN, Hilden, Germany) following the manufacturer’s protocol. In six cases of OCCC, five tumor sections that were cut at least 1 cm apart were sampled with macrodissection. In seven cases of OCCC, five areas of the tumor epithelial component and three areas of the tumor stromal component were sampled with LCM (described below) from one section. In endometriosis and eutopic endometrium, hematoxylin and eosin staining was used to confirm the presence of endometriotic epithelium or endometrial glandular epithelium. FFPE samples were sliced to a thickness of 10 μm and placed on a glass slide with foil (Leica, Wetzlar, Germany), stained with toluidine blue (see supplementary material, Table S1), and cut out by LCM using a Leica LMD7000 (Leica, Wetzlar, Germany) (Figure 1A–E). For each case, 4–12 eutopic endometrial glandular epithelium samples were sampled individually, and 3–4 endometrial stroma in close proximity were grouped together to form one sample. DNA was extracted from the collected samples using the QIAamp DNA Micro Kit (QIAGEN, Hilden, Germany) according to the manufacturer’s protocol.

**Figure 1.**
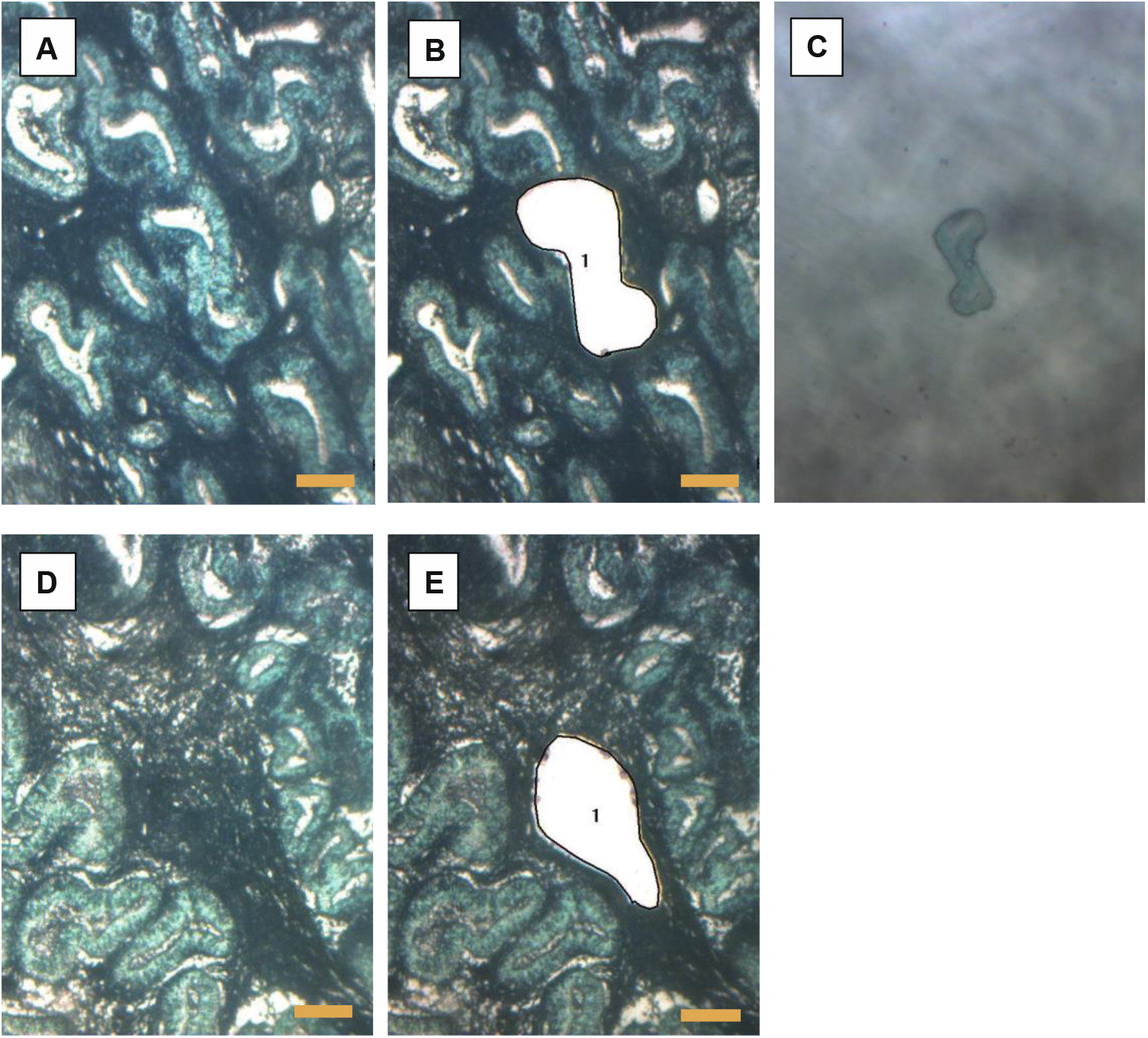
Examples of laser capture microdissection. A and B, Endometrial glands of secretory phase (magnification, 63×). C, Collected endometrial glands. D and E, Endometrial stroma of secretory phase (63×). Scale bars: 100 µm.

### ddPCR assays

Reactions were prepared using 1.1 μL of PrimePCR for ddPCR *PIK3CA* E542K, E545K, or H1047R (Bio-Rad Laboratories, Hercules, CA, USA), 11 μL of ddPCR supermix for probes (no dUTP) (Bio-Rad Laboratories), 5.9 μL of distilled water, and 4 μL of extracted DNA for a total of 22 μL. Droplets were prepared using the Automated Droplet Generator (Bio-Rad Laboratories). After amplification in a thermal cycler (see supplementary material, Table S2), the numbers of wild-type and mutant copies per 20 μL were counted using the QX200 Droplet Digital PCR System (Bio-Rad Laboratories). The mutant allele frequency (MAF) was defined as the ratio of the number of mutant copies to the total number of copies for each hotspot. If any one of the three hotspots (*PIK3CA* E542K, E545K, or H1047R) was not detected in either wild type or mutant copies, that sample was excluded from the analysis.

The cut-off of MAF was set at 15% based on the previous report [7]. Samples with a MAF ≥15% of *PIK3CA* are shown in Figure 2A. The heat map of MAF without setting the cut-off value is shown in Figure 3C and supplementary material, Figure S1.

**Figure 2.**
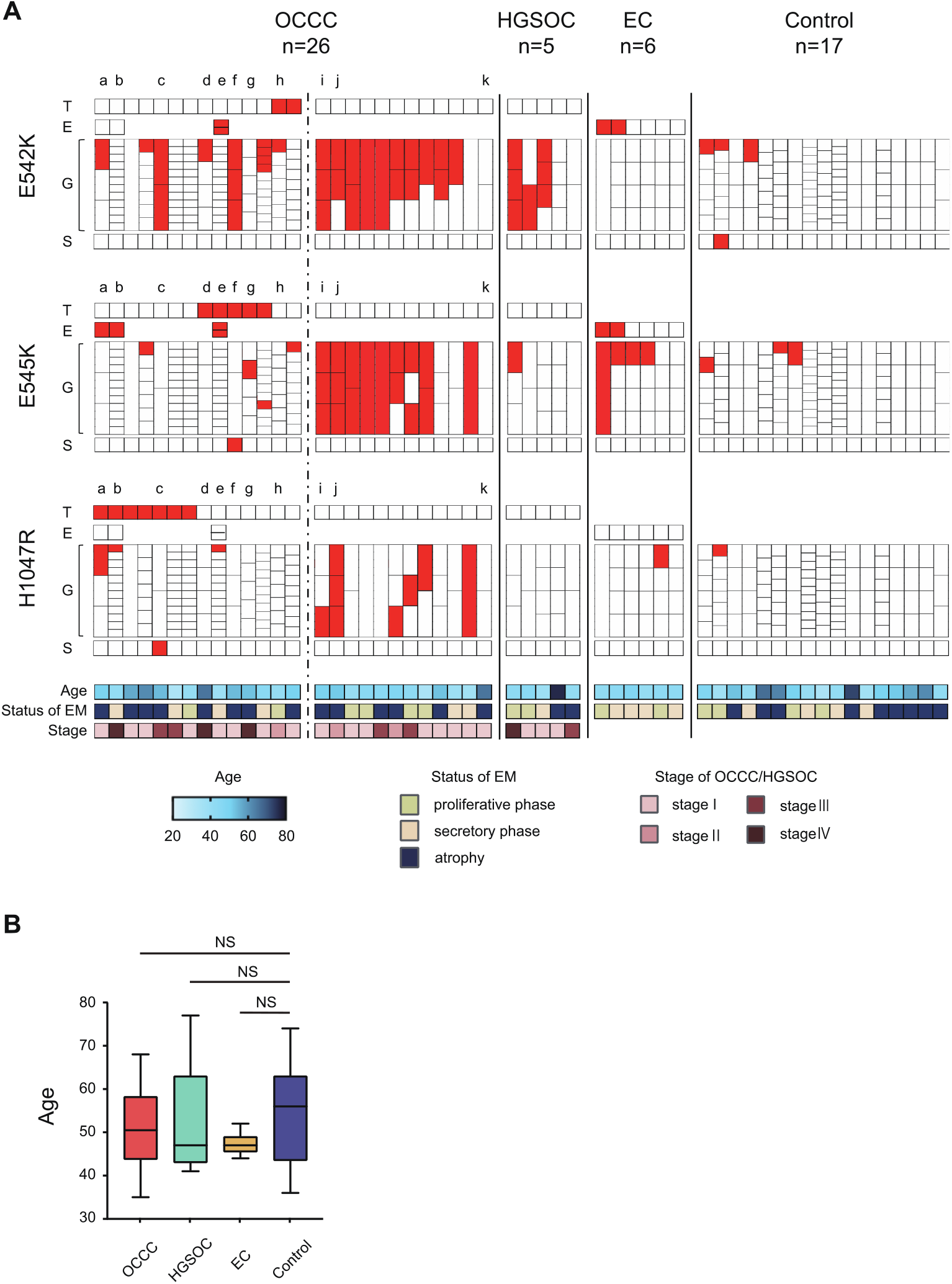
*PIK3CA* mutations in OCCC, HGSOC, EC, endometrial glands, and endometrial stroma. A, *PIK3CA* mutation in OCCC, HGSOC, EC, endometrial glands, and endometrial stroma in all cases. Tumor, endometriosis, endometrial glands, and endometrial stroma of each case were divided into three hot spots and arranged by sample. The spots with mutant allele frequency ≥15% are shown in red. For the OCCC cases, the cases on the left side of the dotted line have *PIK3CA* mutation in the tumor, while the cases on the right side are cases without *PIK3CA* mutation. a–k show cases which tumors were macro- or microdissected and multi-sampled (details are shown in Fig. 3C). B, Comparison of the patient age of each group. The box and whisker plot shows the age of each group. OCCC: ovarian clear cell carcinoma, HGSOC: high-grade serous ovarian carcinoma, EC: endometriotic cyst, T: tumor, E: endometriosis, G: endometrial gland, S: endometrial stroma, EM: endometrium, NS: not significant.

**Figure 3.**
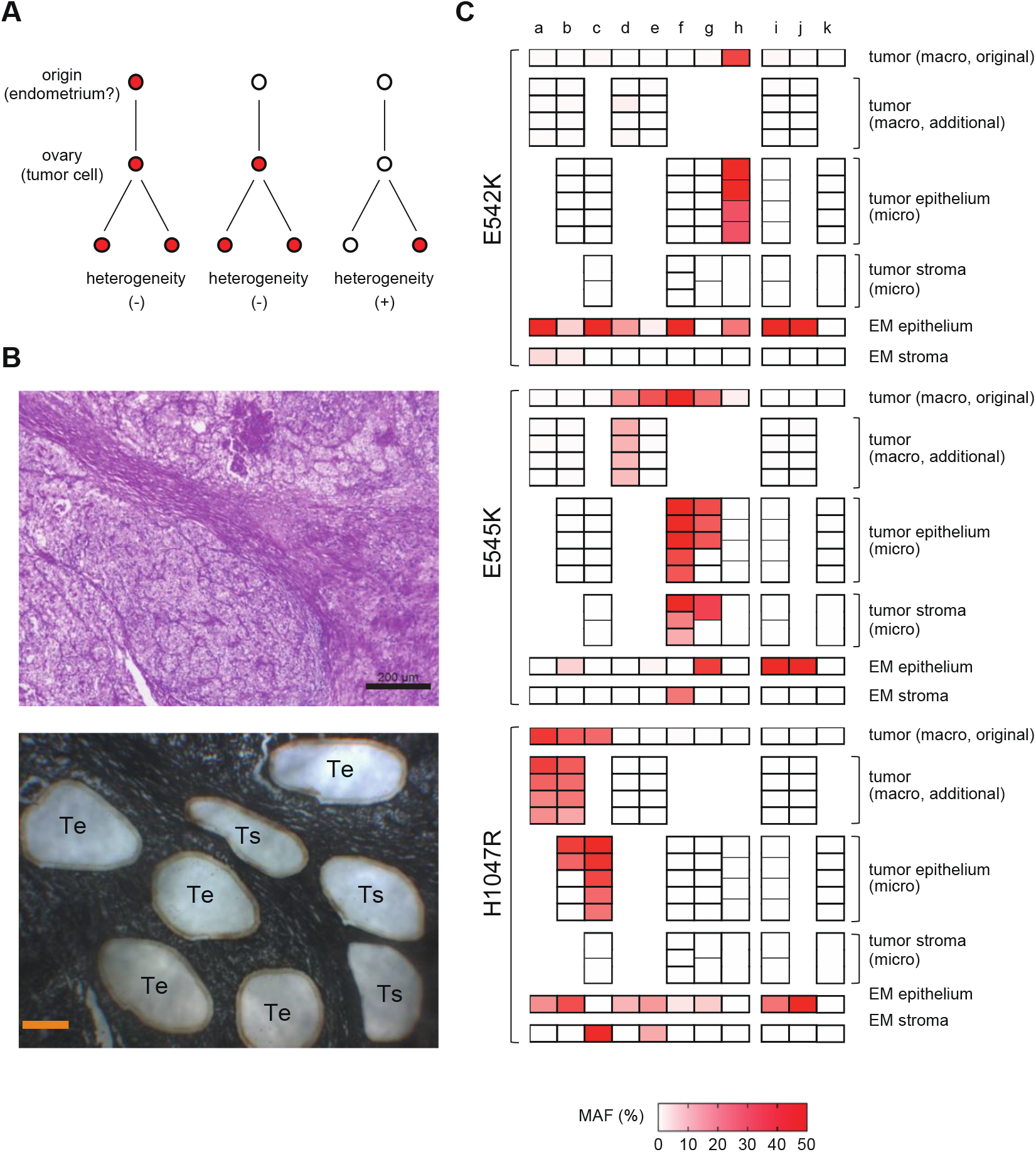
*PIK3CA* mutations in tumors by macro- and microdissection. A, Association between *PIK3CA* mutations in tumor cells of origin and intratumor heterogeneity. If cells with driver *PIK3CA* mutations arise outside of the ovary, such as in the endometrium, and are transported to the ovary to form a tumor (left) or if the tumor cells have driver *PIK3CA* mutations at the time they arise in the ovary (middle), then *PIK3CA* mutations are likely to be present in all tumor cells. However, if *PIK3CA* mutations are not present at the time the tumor cells arise and emerge later, intratumor heterogeneity of *PIK3CA* mutations would occur (right). B, Example of laser capture microdissection. Clear cell carcinoma (case g from Fig. 2, hematoxylin and eosin staining; magnification 63×, toluidine blue staining; magnification 50×). ‘Te’ represents the epithelial component of the tumor and ‘Ts’ represents the stromal component of the tumor. Scale bar: 200 µm. C, MAF of the epithelial component and stromal component of the tumor and eutopic endometrial glandular epithelium and stroma. MAFs of *PIK3CA* by macro- and microdissection are shown in the heat map. For the endometrial glandular epithelium, the sample with the highest MAF was extracted. The cases a-k in B and C are the same as those shown in Fig. 2A. The tumor of cases a, b, d, e, i, j were multiple macrodissected. The epithelial and stromal components of the tumor of cases b, c, f, g, h, i, k were microdissected. Macro: macrodissection; micro, laser-capture microdissection; EM, endometrium; MAF, mutant allele frequency.

### Analysis of public data

A previous report examined endometrial cancer and ovarian cancer cases using Pap smear and plasma, and we downloaded the supplemental target sequencing data for each available specimen [18]. We selected ovarian cancer cases in which *PIK3CA* mutation was detected in the tumor, Pap smear, or plasma and examined the relationship between mutations.

### Statistical analysis

Statistical analyses were performed using the GraphPad Prism ver. 9.0.0 (GraphPad Software, San Diego, CA, USA). Fisher’s exact test was used to compare the proportions between groups, and Mann-Whitney test was used to compare the MAF. The Kaplan-Meier method was used to calculate overall survival, and the log-rank test was used to compare the curves. The correlation between age and MAF was analyzed by Spearman’s rank correlation analysis, and *P* values less than 0.05 were considered statistically significant.

## Results

### *PIK3CA* mutations in ovarian cancer and endometriotic epithelium

We initially obtained FFPE samples for OCCC tumors from 60 cases. The mean age of the patients was 53.9±10.0 years, with 70% of patients in FIGO stage 1 or 2 and 30% in stage 3 or 4. When the cutoff value of MAF was set at 15%, 15 of the 60 tumors (25%) had *PIK3CA* E542K, E545K, or H1047R mutation and one tumor had MAF >5% (E542K, MAF=5.5%). In these 16 cases, *PIK3CA* E542K, E545K, and H1047R mutations in the tumor were mutually exclusive. We divided the 60 patients into two groups according to the presence of *PIK3CA* mutation in the tumor and found no significant difference in overall survival between the two groups (see supplementary material, Figure S2A). There was also no significant difference in age between the two groups (see supplementary material, Figure S2B).

Among the 16 cases of OCCC with *PIK3CA* mutation in the tumor, 14 cases had analyzable eutopic endometrium. In three of these cases, four FFPE samples containing EC or DIE not adjacent to the tumor were available. All three cases showed *PIK3CA* mutation in the endometriotic epithelium. Two of the cases showed E545K mutation in the endometriotic epithelium with H1047R mutation in the tumor (Figure 2A; see supplementary material, Figure S1; case a, b). In the other case, E545K and E542K mutations occurred simultaneously in the endometriotic epithelium with E545K mutation in the tumor (Figure 2A; see supplementary material, Figure S1; case e).

For comparison with the 14 OCCC cases with *PIK3CA* mutation in the tumor, we obtained FFPE samples of tumor and eutopic endometrium from 12 OCCC cases without *PIK3CA* mutation in the tumor, 5 HGSOC cases that were unrelated to endometriosis, and 6 EC cases. In addition, FFPE samples of control endometrium from 17 healthy controls were obtained. The characteristics of the OCCC, HGSOC, EC, and control groups are shown in Table 1. There was no significant difference in age between the groups (Figure 2B). There were no cases of HGSOC with *PIK3CA* mutation with MAF ≥1% in the tumor (Figure 2A; see supplementary material, Figure S1). Endometriotic epithelium of two of the six EC cases showed simultaneous MAF ≥15% mutation in E542K and E545K (Figure 2A).

**Table 1.** Caracteristics of patients

|  | OCCC | HGSOC | EC | Control |
| --- | --- | --- | --- | --- |
| n | 26 | 5 | 6 | 17 |
| Age (mean) | 51.3±9.5 | 51.8±14.0 | 47.3±2.7 | 53.4±11.0 |
| Disease |  |  |  |  |
| stage I | 16 | 3 | - | - |
| stage II | 3 | 0 | - | - |
| stage III | 4 | 1 | - | - |
| stage IV | 3 | 1 | - | - |
| cervical intraepithelial neoplasia | - | - | - | 14 |
| benign ovarian tumor | - | - | - | 3 |
| Status of endometrium |  |  |  |  |
| proliferative phase | 6 | 2 | 2 | 4 |
| secretory phase | 6 | 1 | 4 | 4 |
| atrophy | 14 | 2 | 0 | 9 |

OCCC; ovarian clear cell carcinoma, HGSOC; high-grade serous ovarian carcinoma, EC; endometriotic cyst

### *PIK3CA* mutations in eutopic endometrial glandular epithelium and stroma and comparisons with tumors

In the overall 54 cases (OCCC, HGSOC, EC, and controls), ddPCR was performed on 374 endometrial glands; 1122 hotspots. 265 endometrial glands; 795 hotspots (71%) for which MAF could be calculated at all three hotspots were included in the analysis. The number of cases with endometrial glands with MAF ≥15% in E542K, E545K, or H1047R was significantly higher in the OCCC with *PIK3CA* mutation group and the OCCC without *PIK3CA* mutation group compared with the control group (Table 2). The number of hotspots with MAF ≥15% was significantly higher in the OCCC with *PIK3CA* mutation group, the OCCC without *PIK3CA* mutation group, the HGSOC group, and the EC group compared with the control group (Table 2). In comparing the distribution of MAF of all endometrial glands within each group, the MAF was significantly higher in the OCCC with *PIK3CA* mutation group, the OCCC without *PIK3CA* mutation group, the HGSOC group, and the EC group compared with the control group (Table 2). Interestingly, there were some cases with two or three hotspots of MAF ≥15% mutations among the three hotspots in the same gland, which were particularly common in the OCCC without *PIK3CA* mutation group (Table 2). There was no correlation between age and MAF of *PIK3CA* mutation in endometrial glands (see supplementary material, Figure S3). The MAF of *PIK3CA* mutation in endometrial glands did not correlate with endometrial status (proliferative phase, secretory phase, or atrophy), nor did it correlate with FIGO stage in ovarian cancer cases (data not shown).

**Table 2.** *PIK3CA* mutations in eutopic endometrial glands

| Group | Cases with mutations (MAF≥15%)<br>in endometrial glands | | | Number of hotspots with mutations<br>(MAF≥ 15%) | | | Mean MAF(%) of<br>endometrial glands | | Cases with multiple hotspot mutations<br>in the same glands | | |
| --- | --- | --- | --- | --- | --- | --- | --- | --- | --- | --- | --- |
|  |  | % | P value |  | % | P value | % | P value |  | % | P value |
| OCCC with *PIK3CA* mutation | 11/14 | 79 | 0.01 | 20/300 | 6.7 | 0.01 | 3.1±10.1 | <0.001 | 1/14 | 7.1 | >0.99 |
| OCCC without *PIK3CA* mutation | 11/12 | 92 | 0.002 | 42/78 | 54 | <0.001 | 32.4±35.4 | <0.001 | 7/12 | 58 | 0.003 |
| HGSOC | 3/5 | 60 | 0.31 | 8/45 | 18 | <0.001 | 9.4±17.5 | <0.001 | 1/5 | 20 | 0.41 |
| Endometriotic cyst | 5/6 | 83 | 0.05 | 5/63 | 7.9 | 0.04 | 3.5±11.3 | 0.01 | 0/6 | 0 | >0.99 |
| control | 5/17 | 29 | — | 7/309 | 2.3 | — | 1.3±8.4 | — | 1/17 | 5.9 | — |

OCCC; ovarian clear cell carcinoma, HGSOC; high-grade serous ovarian carcinoma, MAF; mutant allele frequency

We sampled eutopic endometrial stroma from all 54 cases by LCM and *PIK3CA* mutation analysis was performed. Endometrial stroma with MAF ≥15% was detected in only three cases (5.6%); three out of 162 hotspots (1.9%). These three cases included two OCCC cases and one control case (Figure 2A).

Among the 14 cases of OCCC with *PIK3CA* mutation, only three cases (21.4%) had *PIK3CA* mutation with MAF ≥15% in the eutopic endometrial glandular epithelium in a single hotspot coincident with the tumor (Figure 2A; case b, g, h). Conversely, the two OCCC with *PIK3CA* mutation cases showed coincident sites of *PIK3CA* mutations in the tumor and endometrial stroma; one of which had H1047R mutation in the endometrial stroma with H1047R mutation in the tumor and the other had E545K mutation in the endometrial stroma with E545K mutation in the tumor (case c, f).

### Comparison of *PIK3CA* mutation in eutopic endometrium and tumor, and analysis of intratumor heterogeneity

Analysis of the intratumor heterogeneity of *PIK3CA* mutations may help provide insights into the origin of OCCC. For example, if eutopic endometrial cells with *PIK3CA* mutation as an oncogenic driver are the origin of OCCC, then all tumor cells of OCCC would have the *PIK3CA* mutation (Figure 3A). Therefore, in six cases of OCCC, additional macrodissection samplings were performed from four tumor sections more than 1 cm apart from each other. In addition, in seven OCCC cases, samples were collected by LCM from five epithelial components and three stromal components of tumors in very close proximity to each other within the same sample as the original macrodissection (Figure 3B). The additional macrodissection showed that in three of the four OCCC cases with *PIK3CA* mutation in the tumor, the additional sample had the same hotspot mutation as the original sample (Figure 3C; case a, b, d). However, in one case, the additional sample did not have the mutation (Figure 3C; case e). The LCM analysis showed that in some cases, all epithelial components had the same hotspot mutation as the original tumor part (Figure 3C; case c, f, h), while in others, some epithelial components did not have *PIK3CA* mutation (Figure 3C; case b, g). In the majority of the LCM samples, the MAF was higher than in the macrodissection samples (Figure 3C). Among OCCC cases with *PIK3CA* mutation in the tumor, only one case had homogeneous *PIK3CA* mutation in the epithelial component of the tumor and a single hotspot mutation with MAF ≥15%, consistent with the tumor, in the eutopic endometrial glandular epithelium (Figure 3C; case h). Conversely, in two cases with *PIK3CA* mutations with MAF ≥15% in a single hotspot in the eutopic endometrial stroma consistent with the tumor, the *PIK3CA* mutations in the epithelial component of the tumor were homogeneous (Figure 3C; case c, f). In two cases, the stromal component of the tumor showed the same hotspot mutation as the epithelial component of the tumor (Figure 3C; case f, g). In OCCC cases without *PIK3CA* mutation in the original sample of the tumor, neither the additional macrodissection sample nor the LCM sample had *PIK3CA* mutation (Figure 3C; case i, j, k).

### Analysis of previously reported public data

We searched for data on *PIK3CA* mutations in non-cancerous sites in ovarian cancer from previous reports. Wang et al. performed genetic mutation analysis in ovarian cancer and endometrial cancer using Pap smear samples, Tao brush samples, and plasma sample-derived cell-free DNA [18]. The authors reported that cancer-associated mutations were found in ovarian cancer cases with a sensitivity of 63% and a specificity of 100%. Therefore, we compared *PIK3CA* mutations in tumors and non-tumor samples in the publicly available ovarian cancer data from the study. Since the proportion of cases in which Tao brush samples were obtained was very small, Pap smear samples and plasma samples were examined.

In Wang’s cohort, 159 cases of HGSOC were included among 201 cases of ovarian cancer, and tumor *PIK3CA* mutations were found in only two cases (1.2%). In ovarian cancer cases, 25 cases showed *PIK3CA* mutation in at least a tumor, Pap smear, or plasma sample. Only 6 of 25 (24%) cases had the same *PIK3CA* mutation in the tumor and Pap smear or plasma (see supplementary material, Figure S4). In 10 cases in which no *PIK3CA* mutation was found in the tumor (8 HGSOC, 1 endometrioid carcinoma, and 1 mucinous carcinoma), *PIK3CA* mutation was detected only in the Pap smear or plasma sample (see supplementary material, Figure S4).

## Discussion

This study was conducted to examine our hypothesis that EAOC originates from eutopic endometrium [9] and determine whether eutopic endometrium has a high frequency of mutations identical to EAOC. However, the results obtained were quite different from our predictions.

In this study, we focused on three hotspots of *PIK3CA*, E542K, E545K, and H1047R, which are frequent in OCCC, using FFPE samples used in clinical practice to ensure a sufficient number of cases. The frequency of *PIK3CA* mutations in OCCC was reported to be approximately 30%-50% [16, 19-21]. In this study, the number of cases with *PIK3CA* mutations with MAF ≥15% was 25%, which was slightly lower than the previously reported rate, but we consider this to be a reasonable frequency because only three hotspots were studied, and it may reflect the general OCCC population. The *PIK3CA* mutations in three hotspots in the tumor were mutually exclusive (Figures 2A, 3C), which is a natural result as oncogenic driver mutation [22, 23].

In contrast to our findings in OCCC, eutopic endometrial glandular epithelium showed an unexpectedly high frequency of *PIK3CA* mutations. In a previous report that examined cancer-associated mutations in histologically normal endometrium by macrodissection, MAF was at most 10% [4]. Even in a report on cancer-associated mutations in DIE, MAF was at most 20% [6]. Conversely, when endometrial glands are selectively sampled, MAF is often higher than 30% [5]. In our present study, there were many endometrial glands with MAF >30% (Figure 2A), which is consistent with the report. The most impressive data we obtained in this study was the high frequency of eutopic endometrial glandular epithelium-specific *PIK3CA* mutations in OCCC cases without *PIK3CA* mutations in the tumor. Interestingly, mutations in two or three of the three hotspots were often observed in the same gland (Figure 2A, Table 2). Recent reports have shown that cancer-associated mutations are frequently found in many normal tissues such as esophagus, skin, trachea, and colon, as well as the endometrium [24-29]. In esophageal cancer cases, the frequency of mutations in *NOTCH1*, a cancer-associated gene, was significantly higher in the normal epithelium of the esophagus adjacent to the cancer than in the esophageal cancer (66% vs. 15%), and there were multiple mutations in *NOTCH1* in the same clone of normal esophageal epithelium [24]. In addition, meticulous analysis of normal esophageal epithelium did not show a lineage to esophageal cancer. These results seem to indicate a similar phenomenon to our data. Furthermore, if *PIK3CA* mutations in eutopic endometrium are driver mutations of OCCC, there would be no heterogeneity of *PIK3CA* mutations in tumors (Figure 3A). However, intratumor heterogeneity of *PIK3CA* mutations was observed in three of the eight cases in which multi-sampling of the tumor was performed, and only one case showed both *PIK3CA* mutation in the eutopic endometrial glandular epithelium and homogeneous *PIK3CA* mutation in the tumor (Figure 3C). These results suggest that *PIK3CA* mutations in eutopic endometrial glandular epithelium are likely to be passenger mutations rather than driver mutations in the genesis of OCCC.

Similar to the results in eutopic endometrium, *PIK3CA* mutations in the endometriotic epithelium coexisting with OCCC were not identical to those in the tumor. Previous studies reported that in EAOC, the same mutations found in cancer were also found in endometriotic lesions adjacent to the cancer [19,30]. However, in our study, we isolated and examined endometriotic epithelium at a site remote from the cancer by LCM, which is different from the method used in previous reports. In endometriotic epithelium, multiple hotspot mutations were also found in the same sample (Figure 2A). Therefore, *PIK3CA* mutations in endometriotic epithelium as well eutopic endometrial glandular epithelium may be passenger mutations.

There are few *PIK3CA* mutations in HGSOC that are not associated with endometriosis [31], and no *PIK3CA* mutations were found in HGSOC tumors in our study. However, *PIK3CA* mutations were found in the eutopic endometrial glands of HGSOC at a higher frequency than in the endometrial glands of controls (Figure 2A, Table 2). In addition, analysis of the data by Wang et al. showed the presence of *PIK3CA* mutations in cell-free DNA of Pap smear and plasma in HGSOC cases (see supplementary material, Figure S4). In a previous report on genome sequencing in normal uterine cervix, *PIK3CA* mutations were not detected at all [32]. The reason for the presence of *PIK3CA* mutations in Pap smear samples from ovarian cancer cases without *PIK3CA* mutations in the tumor may be that *PIK3CA* mutations are more likely to occur in the cervix in ovarian cancer cases. Alternatively, it is also possible that the *PIK3CA* mutation occurred in the eutopic endometrial glands, as we observed, and DNA or endometrial cells with *PIK3CA* mutation were introduced into the Pap smear samples. In ovarian cancer, cell-free DNA in plasma and tumor mutations do not always coincide, and intratumor heterogeneity has been commonly considered as a cause of this phenomenon [33]. Our data, however, indicate that other genetic variants of normal organ origin must be kept in mind. Individuals with a history of cancer, including OCCC, have a higher risk of developing a second cancer than individuals without a history of cancer [34-36]. This may be because the carcinogenic stress that caused the tumor in patients with carcinoma may also affect organs unrelated to the tumor, causing the accumulation of genetic variants in the other organs. Alternatively, considering the frequency with which humans develop cancer during their lifetime, many cancer-associated mutations that occur in normal tissues may be passenger mutations. The biological significance of cancer-associated mutations in normal tissues should be investigated in future studies.

Surprisingly, there were some cases in which the stromal component of OCCC had *PIK3CA* mutations, and these MAFs were as high as those of the epithelial component, which did not appear to be contamination (Figure 3C). This suggests that the stromal component of OCCC also contains tumor cells. Previous studies showed that the side-population cells of endometrial cancer can differentiate into mesenchymal cells [37], and OCCC tumor cells may also exist as mesenchymal cells. Among the OCCC cases in this study, two had *PIK3CA* mutations with MAF ≥15% in the eutopic endometrial stroma, and both had the same hotspot mutation in the tumor (Figure 2A). Moreover, in both cases, microdissection revealed identical hotspot mutations in all samples of the epithelial component of the tumor, and one showed homogeneous identical hotspot mutations not only in the epithelial component but also in the stromal component of the tumor (Figure 3). The eutopic endometrial stroma has been reported to contain few cancer-associated mutations [38]. However, our data suggest that low frequency mutations in endometrial stromal cells contribute more to the true driver mutations of EAOC than high frequency, probably mostly passenger, cancer-associated mutations in endometrial glandular epithelial cells, and eutopic endometrial stromal cells may be the origin of EAOC. The fact that endometrial stromal cells have been used as material in many studies on endometriosis may also support our findings [39]. Our data, however, are based on a very limited number of cases and do not explain the process of transformation from stromal cells to epithelial cells, so we have only shown one possibility. Nevertheless, our results suggest that future studies of the eutopic endometrium for the purpose of searching for the origin of EAOC should examine not only the endometrial glandular epithelium but also the endometrial stroma.

In this study, hotspot mutations of *PIK3CA* were examined by ddPCR using FFPE samples because of the ease of sample collection. However, this method did not allow us to conclude whether the origin of OCCC was eutopic endometrium. For future research purposes, it will be necessary to cryopreserve the removed specimens, including the normal-looking uterus, when performing surgery for ovarian cancer. A large number of endometrial glands and stroma from the entire endometrium should then be collected by LCM and analyzed by whole exome or whole genome sequencing and compared with the data from tumor tissue. Further advances in sequencing technology are therefore needed for these studies.

In conclusion, here we found that *PIK3CA* mutations specific to endometrial glandular epithelium are frequently observed in eutopic endometrium with OCCC, and most of the mutations are passenger mutations that are not related to *PIK3CA* mutations in OCCC. Further comprehensive analysis is needed to clarify the significance of cancer-associated mutations in eutopic endometrium.

## Acknowledgements

We thank Akiko Kyoda for FFPE sample preparation and toluidine blue staining. This study was supported in part by Japan Society for the Promotion of Science (JSPS) KAKENHI grant number 18H02947 (Grant-in-Aid for Scientific Research B for Noriomi Matsumura) and 20K21665 (Grant-in-Aid for Challenging Exploratory Research for Noriomi Matsumura).

## Authors’ Contributions

KM and NM designed the study. AK, HN and YK collected specimens for the study, KM and CM performed experiments. KS and HT performed data analysis. KM drafted the manuscript. KN and NM supervised the study. All authors were involved in writing the paper and had final approval of the submitted and published versions.

