## Supplementary Figure S1 for "Frequent *PIK3CA* mutations in eutopic endometrium of patients with ovarian clear cell carcinoma"

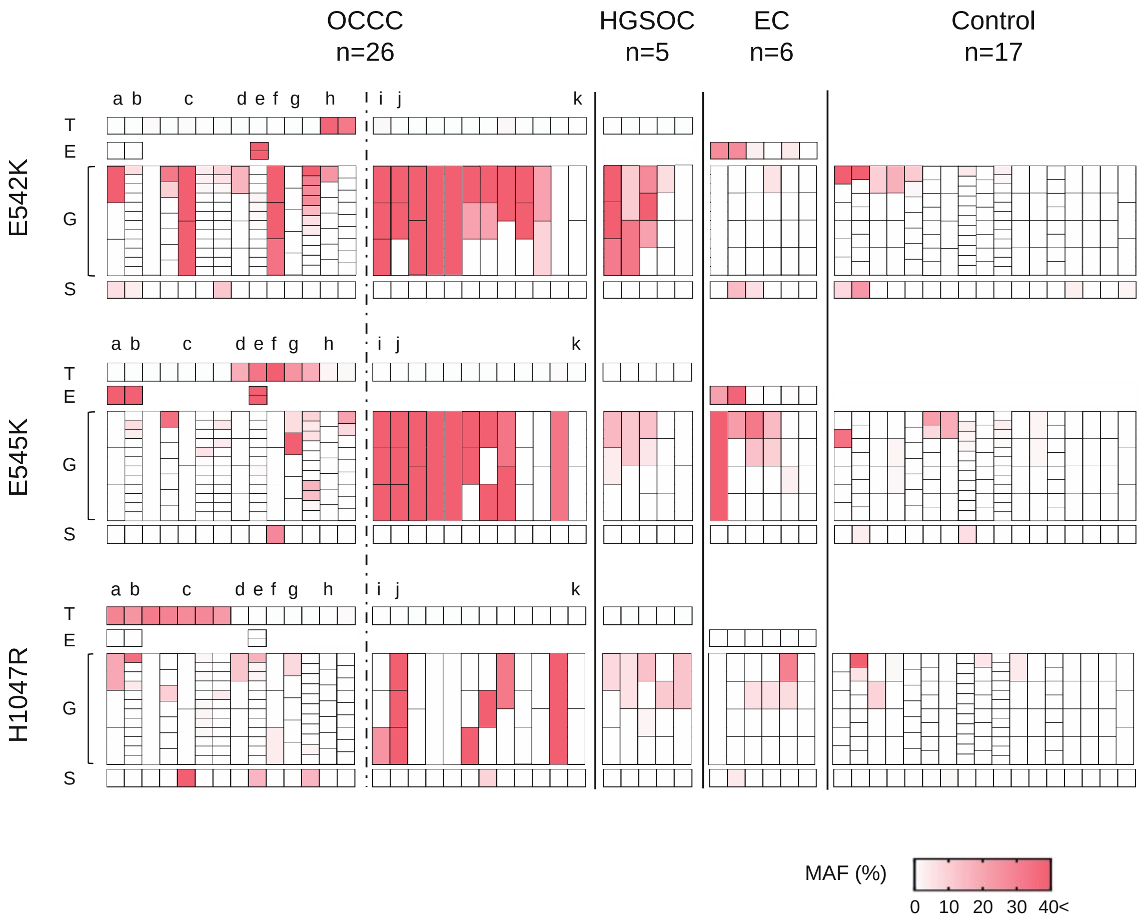


**Supplementary Figure S1.** Heatmap of the MAF of *PIK3CA* in OCCC, endometriosis, and endometrium in all cases. The samples are arranged as in Fig. 2A, and all MAF values are shown in the heat map. For the OCCC cases, the cases on the left side of the dotted line have *PIK3CA* mutation in the tumor, while the cases on the right side are cases without *PIK3CA* mutation. a–k show cases in which tumors were macro- or microdissected and multi-sampled (details are shown in Fig. 3). MAF: mutant allele frequency, OCCC: ovarian clear cell carcinoma, HGSOC: high-grade serous ovarian carcinoma, EC: endometriotic cyst, T: tumor, E: endometriosis, G: endometrial gland, S: endometrial stroma.
