## Supplementary Figure S2 for "Frequent *PIK3CA* mutations in eutopic endometrium of patients with ovarian clear cell carcinoma"

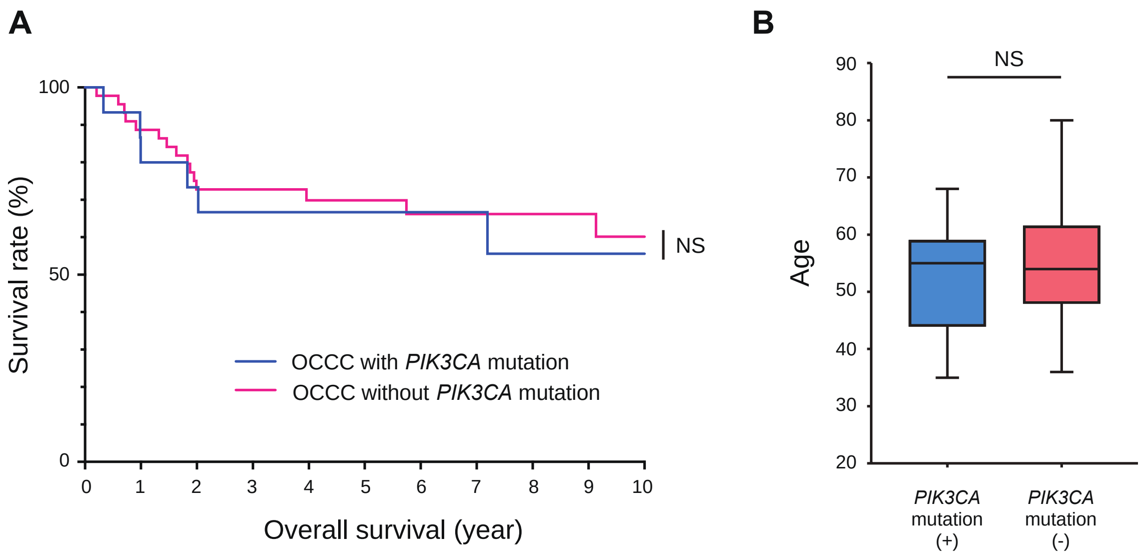


**Supplementary Figure S2.** Survival and age in OCCC cases according to *PIK3CA* mutation. A, Overall survival of 60 OCCC cases. Patients were divided into two groups according to the presence of *PIK3CA* mutation with MAF ≥15%. Overall survival is shown by Kaplan–Meier curve. The blue line indicates the group with MAF ≥15% of *PIK3CA* mutation, and the red line indicates the other group. B, Age. Patients were divided into two groups according to the presence of *PIK3CA* mutation with MAF ≥15%. Age of each group is shown in a box and whisker plot. NS: not significant. OCCC: ovarian clear cell carcinoma.
