## Supplementary Figure S3 for "Frequent *PIK3CA* mutations in eutopic endometrium of patients with ovarian clear cell carcinoma"

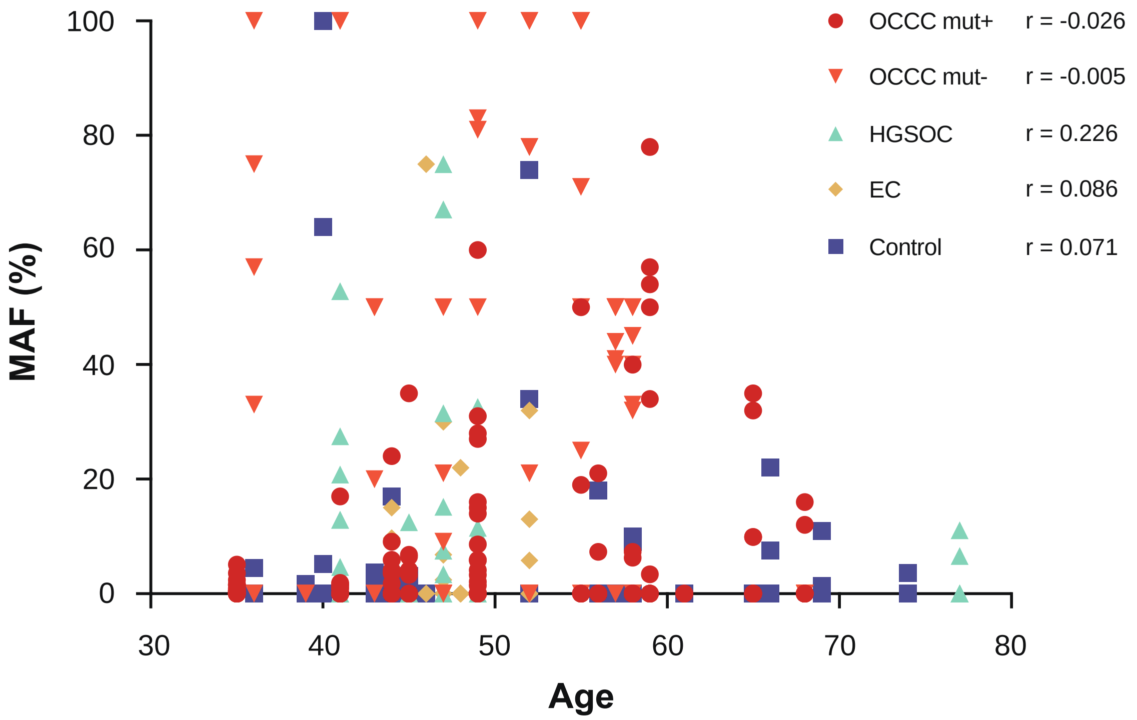


**Supplementary Figure S3.** Correlation between eutopic endometrial glands and age. Spearman's rank correlation coefficient was calculated for age and MAF. There was no correlation between age and MAF in the group with *PIK3CA* mutation with MAF ≥15% in the tumor of OCCC (OCCC mut+), the group without mutation (OCCC mut-), HGSOC, EC, and controls. OCCC: ovarian clear cell carcinoma, HGSOC: high-grade serous ovarian carcinoma, EC: endometriotic cyst, MAF: mutant allele frequency.
