## Supplementary Figure S4 for "Frequent *PIK3CA* mutations in eutopic endometrium of patients with ovarian clear cell carcinoma"

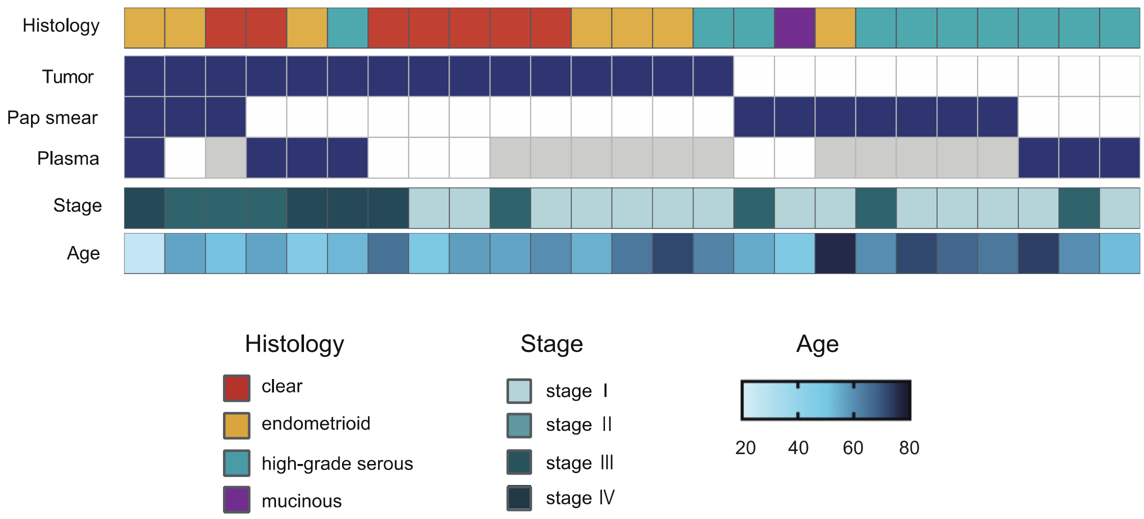


**Supplementary Figure S4.** Twenty-five ovarian cancer cases with *PIK3CA* mutations in tumor, Pap smear, and plasma from the study by Wang et al. Tumor, Pap smear, and plasma with *PIK3CA* mutations are shown in dark blue. Cases that did not have samples are shown in gray.
