## Supplementary Table S1 for "Frequent *PIK3CA* mutations in eutopic endometrium of patients with ovarian clear cell carcinoma"

**Supplementary Table S1** Toluidine blue staining

| |
|---|
| Deparaffinization with xylene |
| ↓ |
| Rinsing with distilled water |
| ↓ |
| Soaking in 0.05% toluidine blue solution for 25 seconds |
| ↓ |
| Remove staining solution and dehydration with anhydrous ethanol |
| ↓ |
| Drying in the air |
