## Supplementary Table S2 for "Frequent *PIK3CA* mutations in eutopic endometrium of patients with ovarian clear cell carcinoma"

**Supplementaly Table S2** Thermal cycles for droplet digital PCR

| Step | Tempature (°C) | Time (sec) | | |
|---|---|---|---|---|
| 1 | 95 | 600 | | |
| 2 | 94 | 30 | * | |
| 3 | 55 | 60 | * | *repated for 39 cycles |
| 4 | 98 | 600 | | |
| 5 | 4 | ∞ | | |
